## Supplemental Figures for "Mapping the immunogenic landscape of near-native HIV-1 envelope trimers in non-human primates"

### Supplemental Data

A

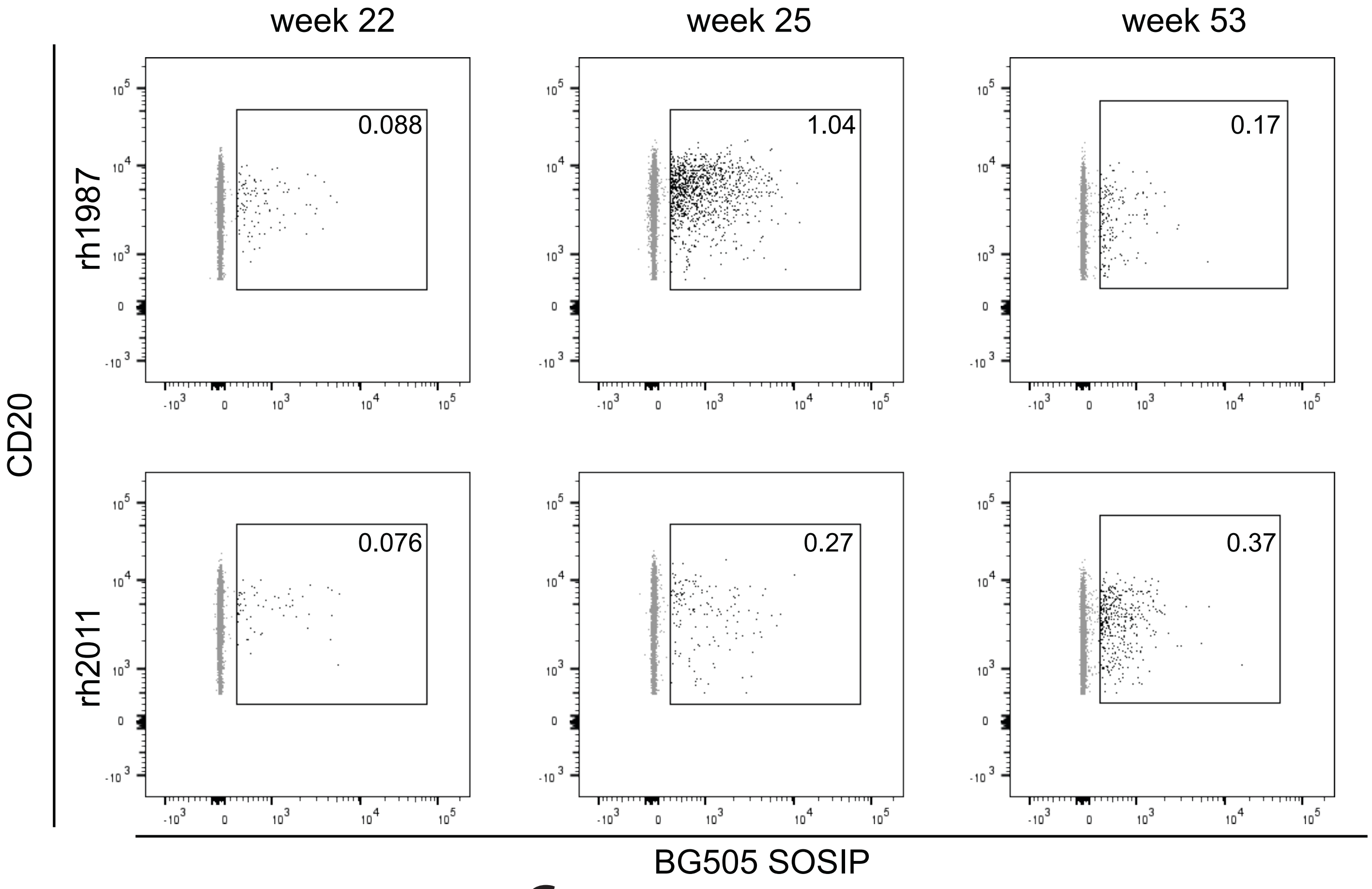


B

| mAb | SOSIP ELISA | gp120 ELISA |
|---|---|---|
| RM19A | ++ | +++ |
| RM19A1 | +++ | +++ |
| RM19A2 | +++ | +++ |
| RM19A3 | ++ | + |
| RM19B | ++ | – |
| RM19B1 | + | – |
| RM19C | + | – |
| RM19C2 | ++ | – |
| RM19C3 | ++ | – |
| RM19C4 | ++ | + |
| RM19D | +++ | +++ |
| RM19E | + | – |
| RM19F | + | – |
| RM19F1 | ++ | – |
| RM19G | ++ | – |
| RM19J | ++ | – |
| RM19K | +++ | ++ |
| RM19L | ++ | – |
| RM19M | ++ | – |
| RM19N | + | + |
| RM19O | +++ | +++ |
| RM19P | +++ | +++ |
| RM19R | ++ | – |
| RM19S | ++ | – |
| RM19T | ++ | ++ |

C

| mAb | SOSIP ELISA | gp120 ELISA |
|---|---|---|
| RM20A | ++ | – |
| RM20A1 | ++ | – |
| RM20A2 | ++ | – |
| RM20A3 | ++ | – |
| RM20B | + | – |
| RM20B1 | + | – |
| RM20C | + | – |
| RM20D | ++ | +++ |
| RM20E | +++ | + |
| RM20E1 | ++ | + |
| RM20E2 | ++ | + |
| RM20E3 | + | + |
| RM20F | ++ | + |
| RM20G | ++ | – |
| RM20H | ++ | ++ |
| RM20I | ++ | ++ |
| RM20J | +++ | +++ |

| EC50 (ug/mL) | |
|---|---|
| <0.1 | +++ |
| 0.1-1 | ++ |
| 1-10 | + |
| >10 | – |

**Figure S1**. MAb isolation and characterization from BG505 SOSIP.664 trimer-immunized macaques. (A) FACS gating strategy for isolation of BG505 SOSIP specific memory B-cells. (B) BG505 SOSIP.664 trimer and BG505 gp120 ELISA binding data for mAbs isolated from RM rh1987. (C) BG505 SOSIP.664 trimer and BG505 gp120 ELISA binding data for mAbs isolated from RM rh2011.

| mAb name | 2D class averages | 3D reconstructions | Epitope | EMDB# |
|---|---|---|---|---|
| RM19A1 | 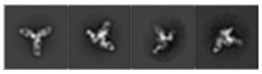 | 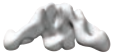 | 289 GH | EMD-21062 |
| RM19B | 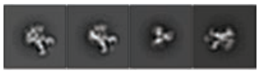 | 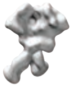 | base | EMD-21075 |
| RM19B1 | 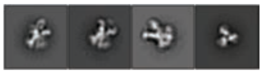 | 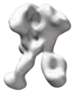 | base | EMD-21077 |
| RM19C | 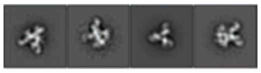 | 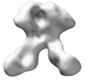 | base | EMD-21078 |
| RM19C2 | 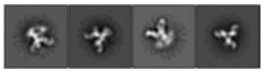 | 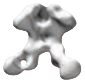 | base | EMD-21079 |
| RM19C3 | 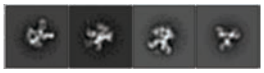 | 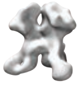 | base | EMD-21082 |
| RM19E | 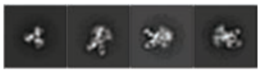 | 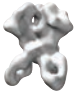 | base | EMD-21080 |
| RM19F | 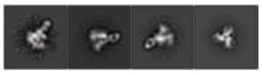 | 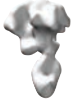 | base | EMD-21056 |
| RM19G | 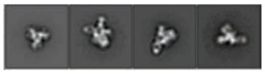 | 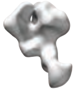 | base | EMD-21081 |
| RM19J | 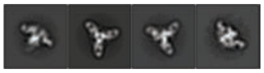 | 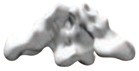 | 289 GH | EMD-21055 |
| RM19K | 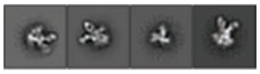 | 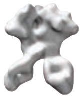 | base | EMD-21076 |
| RM19L | 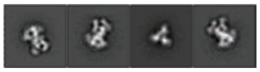 | 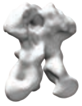 | base | EMD-21066 |
| RM19M | 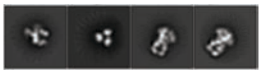 | 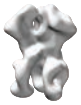 | base | EMD-21061 |
| RM19N | 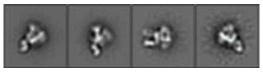 | 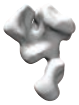 | base | EMD-21053 |
| RM19O | 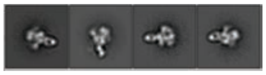 | 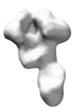 | base | EMD-21065 |
| RM19P | 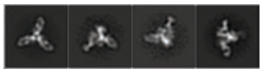 | 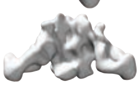 | 289 GH | EMD-21064 |
| RM19R | 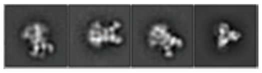 | 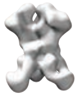 | base | EMD-21058 |
| RM19S | 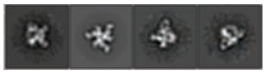 | 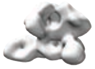 | N611/FP | EMD-21059 |
| RM19T | 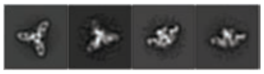 | 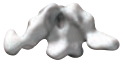 | N289 GH | EMD-21057 |

| mAb name | 2D class averages | 3D reconstructions | Epitope | EMDB# |
|---|---|---|---|---|
| RM20A2 | 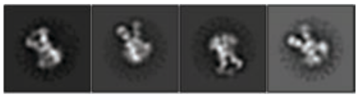 | 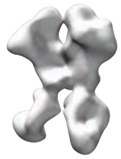 | base | EMD-21083 |
| RM20A3 | 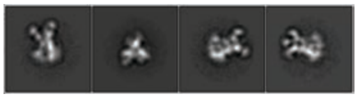 | 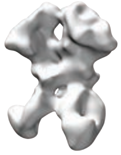 | base | EMD-21084 |
| RM20B | 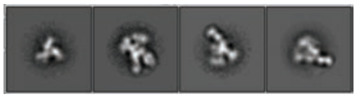 | 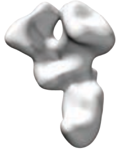 | base | EMD-21085 |
| RM20B1 | 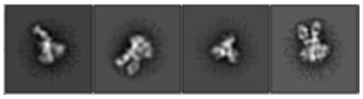 | 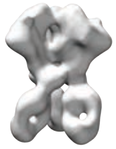 | base | EMD-21086 |
| RM20C | 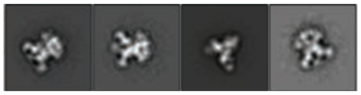 | 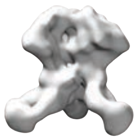 | base | EMD-21087 |
| RM20E | 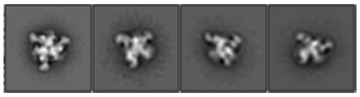 | 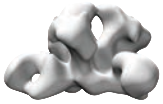 | N611/FP | EMD-21093 |
| RM20E1 | 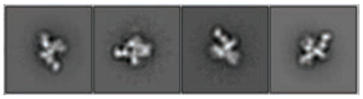 | 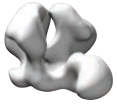 | N611/FP | EMD-21090 |
| RM20F | 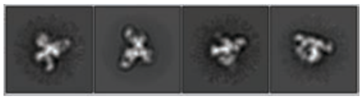 | 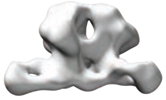 | gp120/gp41 interface | EMD-21091 |
| RM20G | 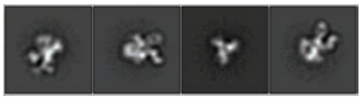 | 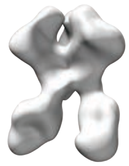 | base | EMD-21088 |
| RM20H | 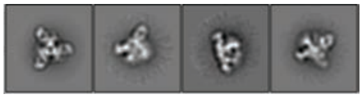 | 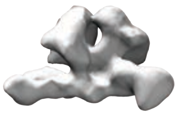 | gp120/gp41 interface | EMD-21092 |
| RM20J | 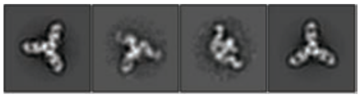 | 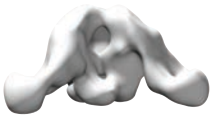 | 289 GH | EMD-21089 |

**Figure S2.** Negative stain electron microscopy epitope mapping of Fabs from rh1987 and rh2011. Representative 2D class averages, 3D reconstructions, and EMDB accession numbers.

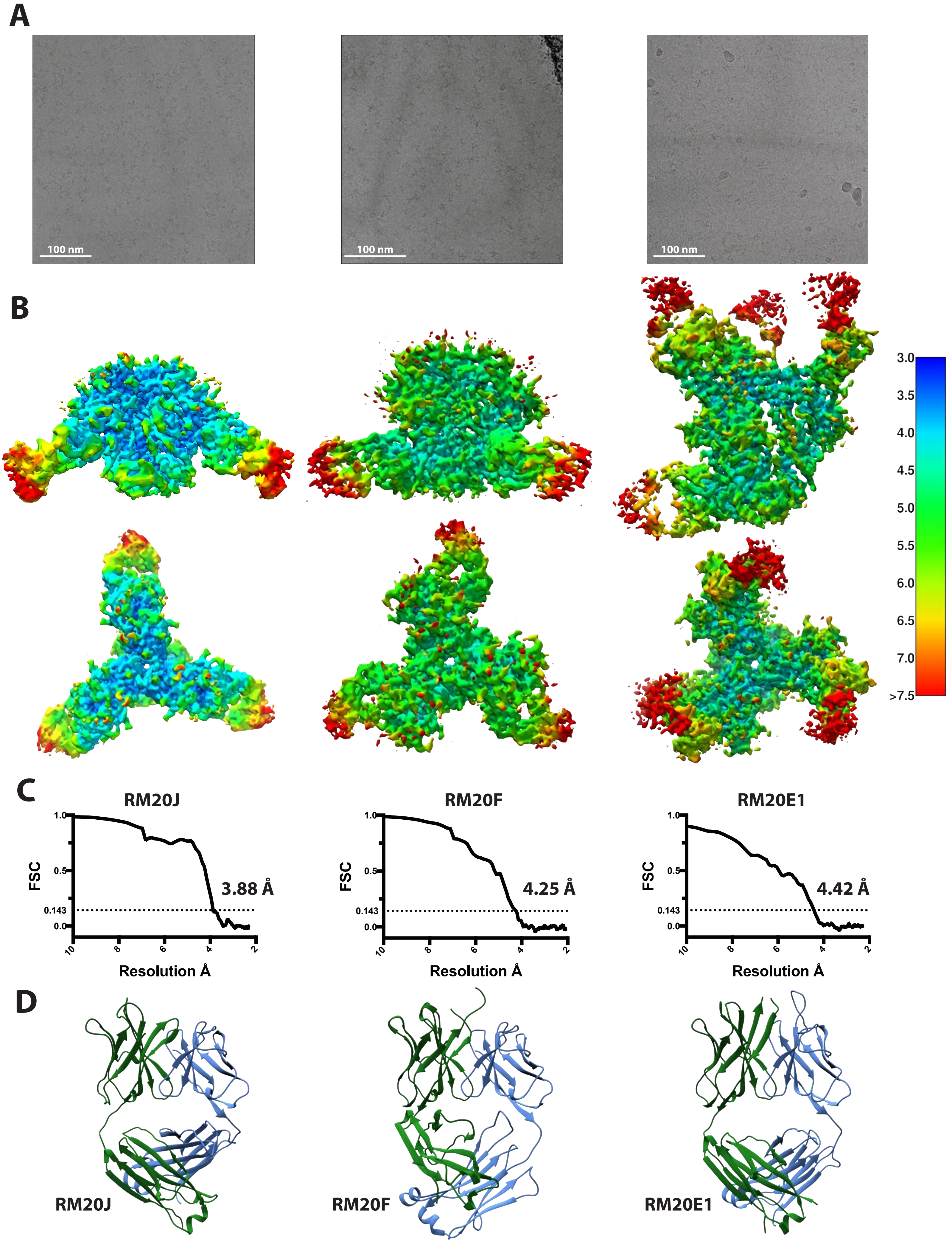

A
100 nm
100 nm
100 nm
B
3.0
3.5
4.0
4.5
5.0
5.5
6.0
6.5
7.0
>7.5
C
RM20J
RM20F
RM20E1
FSC
1.0
0.5
0.143
0.0
3.88 Å
4.25 Å
4.42 Å
10
8
6
4
2
Resolution Å
D
RM20J
RM20F
RM20E1

**Figure S3.** Cryo-EM structures of BG505 SOSIP trimers with mAbs and crystal structures of mAbs**.** (A) Representative micrographs for cryoEM datasets. (B) Local resolution maps for each complex generated in cryoSPARC v2 (Punjani et al., 2017). (C) Gold-standard Fourier shell correlation (FSC) curves for each complex showing global resolution calculated at FSC = 0.143. (D) Crystal structures of unliganded Fabs RM20J, RM20F, and RM20E1. Heavy chains are shown in green and light chains shown in light blue.

| Virus name | Tier* | Subtype | Accession | RM20F | ACS202* | VRC34* |
|---|---|---|---|---|---|---|
| 001428_2_42 | 2 | C | EF117266 | >100 | >10 | 0.075 |
| 1012_11_TC21_3257 | 1B or 2 | B | EU289184 | >100 | >10 | n.d. |
| AC10_29 | 2 | B | AY835446 | >100 | >10 | >10 |
| BJOX010000_06_2 | 2 | 01_AE | HM215373 | >100 | 0.506 | >10 |
| BJOX015000_11_5 | 2 | 01_AE | HM215377 | >100 | 0.023 | n.d. |
| BJOX025000_01_1 | 2 | 01_AE | HM215386 | >100 | 0.003 | >10 |
| BJOX028000_10_3 | 2 | 01_AE | HM215389 | >100 | 0.005 | >10 |
| C2101_C1 | 2 | 01_AE | JN944661 | >100 | n.d. | 0.218 |
| C3347_C11 | 2 | 01_AE | JX512902 | >100 | >10 | >10 |
| C4118_9 | 2 | 01_AE | JQ352782 | >100 | 0.411 | >10 |
| CAP210_E8 | 2 | C | DQ435683 | >100 | 0.297 | 0.183 |
| CAP45_G3 | 2 | C | DQ435682 | >100 | n.d. | 0.056 |
| CNE20 | 2 | 07_BC | HM215406 | >100 | >10 | >10 |
| CNE5 | 2 | 01_AE | HM215415 | >100 | >10 | >10 |
| DU156_12 | 2 | C | DQ411852 | >100 | n.d. | >10 |
| DU172_17 | 2 | C | DQ411853 | >100 | n.d. | 0.078 |
| PVO_4 | 2 or 3 | B | AY835444 | >100 | 0.043 | >10 |
| Q23_17 | 1B | A1 | AF004885 | >100 | >10 | 0.099 |
| Q842_D12 | 2 | A1 | AF407160 | >100 | n.d. | 0.138 |
| REJO4541_67 | 2 | B | AY835449 | >100 | >10 | >10 |
| RHPA4259_7 | 2 | B | AY835447 | >100 | 0.012 | 1.559 |
| SC422_8 | 2 | B | AY835441 | >100 | >10 | >10 |
| THRO4156 | 2 | B | AY835448 | >100 | n.d. | >10 |
| TRJO4551_58 | 3 | B | AY835450 | >100 | 0.029 | 7.334 |
| TRO_11 | 2 | B | AY835445 | >100 | >10 | >10 |
| ZM109_4 | 1B or 2 | C | AY424138 | >100 | n.d. | 1.432 |
| ZM135_10A | 2 | C | AY424079 | >100 | n.d. | >10 |
| ZM197_7 | 1B or 2 | C | DQ388515 | >100 | n.d. | >10 |

**Figure S4.** Neutralization data for a panel of 40 heterologous pseudoviruses against RM20F, ACS202, and VRC34. *Data obtained from Los Alamos National Lab (LANL) CATNAP application (Yoon et al., 2015).

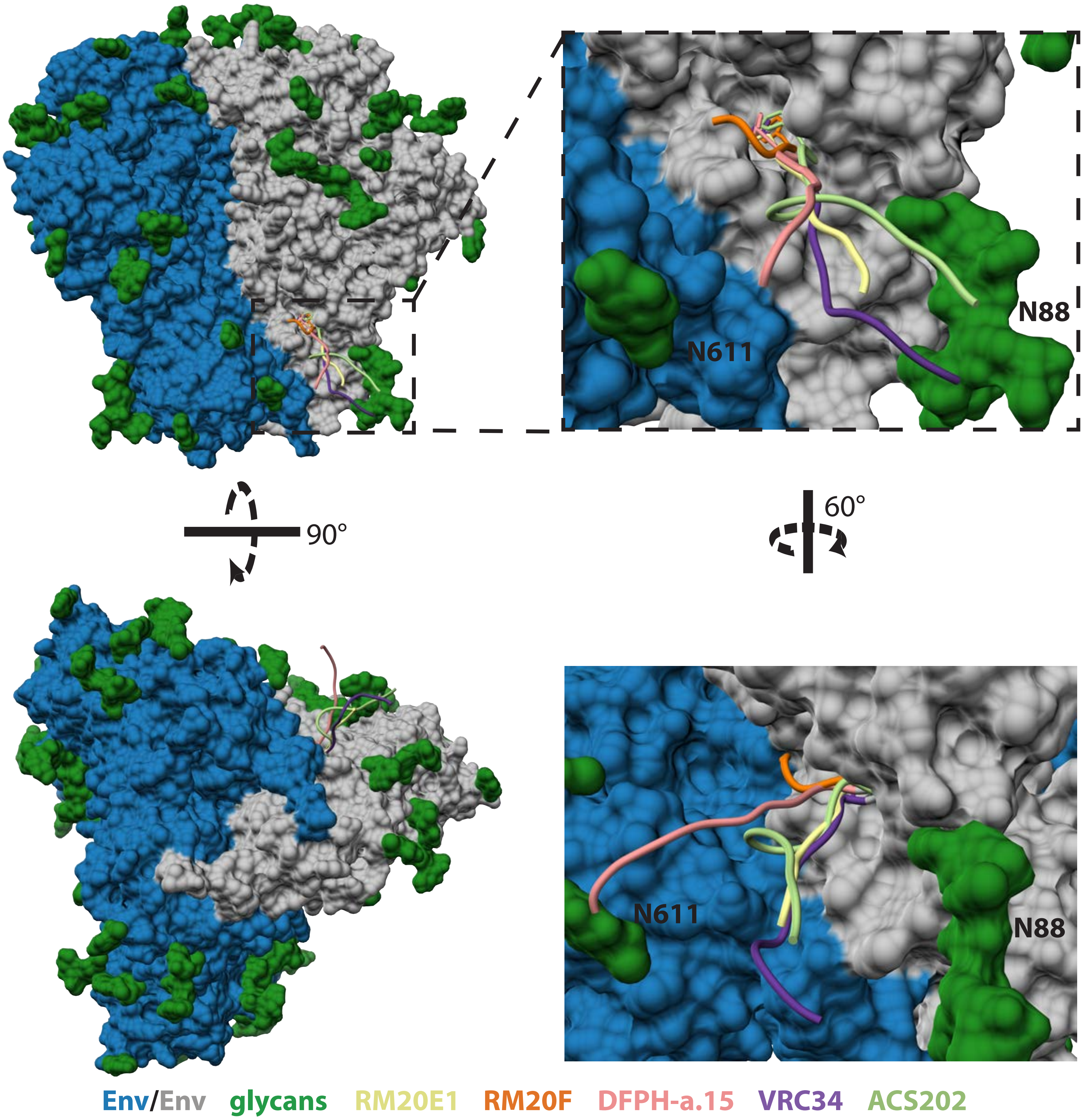


**Figure S5.** Antibody bound FP conformations. The HIV Env trimer is shown as a surface representation with one protomer colored in grey, the other two protomers colored in blue, and the N-linked glycans colored in green. The FP (residues 512-522) from the antibody bound structures of RM20E1, RM20F, DFPH-a.15 (PDB: 6N1W), VRC34 (PDB: 5I8H), and ACS202 (PDB: 6NC2) are shown as colored backbone ribbon diagrams.

#### Supplementary Table 2. MAb Characteristics

| mAb | Timepoint | Animal ID | LC type | VH gene | D gene | JH gene | HCDR3 aa | HCDR3 length | HC %SHM | VL gene | JL gene | LCDR3 aa | LCDR3 Length | LC %SHM | Epitope Targeted |
|---|---|---|---|---|---|---|---|---|---|---|---|---|---|---|---|
| RM19A | week 22 | rh1987 | Kappa | IGHV3-AFE*01_S3736 | IGHD4-22*01 | IGHJ4*01 | ARGGVYSNYEF | 11 | 4.3% | IGKV1-AAL*01_S2543 | IGKJ1-I1 | QQYNSYPPT | 9 | 5.3% | 289 glycan hole* |
| RM19A1 | week 25 | rh1987 | Kappa | IGHV3-AFE*01_S3736 | IGHD4-22*01 | IGHJ4*01 | AKGGVYSNYDF | 11 | 6.3% | IGKV1-AAL*01_S2543 | IGKJ1-I1 | QQYNSDPPT | 9 | 4.4% | 289 glycan hole |
| RM19A2 | week 25 | rh1987 | Kappa | IGHV3-AFE*01_S3736 | IGHD4-22*01 | IGHJ4*01 | AKGGVYSNYDF | 11 | 7.2% | IGKV1-AAL*01_S2543 | IGKJ1-I1 | QQYNSDPPT | 9 | 5.6% | 289 glycan hole* |
| RM19A3 | week 25 | rh1987 | Kappa | IGHV3-AFE*01_S3736 | IGHD4-22*01 | IGHJ4*01 | ARGGVYSNYDY | 11 | 3.4% | IGKV1-AAL*01_S2543 | IGKJ1-I1 | QQHNRDPPT | 9 | 4.0% | 289 glycan hole* |
| RM19D | week 22 | rh1987 | Kappa | LJI.Rh_IGHV3.50 | IGHD3-26*01 | IGHJ6*01 | VRDYYGGGLD | 10 | 3.3% | IGKV2-AEB*01 | IGKJ2-I2 | TQGLEFPYS | 9 | 2.6% | n.d. |
| RM19J | week 25 | rh1987 | Kappa | IGHV4-AGU*01_S6650 | LJI.Rh_IGHD4.24 | IGHJ4*01 | ARFNSGDYDF | 10 | 8.6% | IGKV1-AAL*01_S2543 | IGKJ3-I3 | HQHNTYPLT | 9 | 5.6% | 289 glycan hole |
| RM19N | week 25 | rh1987 | Kappa | IGHV4-S40_S2532 | IGHD2-8*01 | IGHJ4*01 | ARERVVSATSDVYYFDY | 17 | 2.1% | IGKV1-S57 | IGKJ2-I2 | QHSYDTPYS | 9 | 4.4% | base |
| RM19P | week 25 | rh1987 | Kappa | IGHV4-AGU*01_S6650 | IGHD6-29*01 | IGHJ4*01 | SSSGWSV | 7 | 7.7% | IGKV1-AAY*01 | IGKJ2-I2 | QQYNSYPYS | 9 | 3.7% | 289 glycan hole |
| RM19R | week 25 | rh1987 | Kappa | IGHV4-S40_S2532 | IGHD3-3*01 | IGHJ5-1*01 | AFFWSTYYKRFDV | 13 | 8.8% | IGKV3-AAP*01_S6980 | IGKJ1-I1 | HQENDWPWT | 9 | 6.0% | base |
| RM19S | week 53 | rh1987 | Kappa | IGHV4-ABB*01_S8200 | IGHD3-9*01 | IGHJ4*01 | ARRSRRYGDDYGYWYSRSFYFDS | 23 | 7.4% | LJI.Rh_IGKV1.56 | IGKJ4-I4 | QQDYSTPFT | 9 | 5.3% | N611/FP |
| RM19T | week 25 | rh1987 | Kappa | IGHV4-AGU*01_S6650 | LJI.Rh_IGHD4.24 | IGHJ4*01 | ARFNSDYDF | 9 | 9.5% | IGKV1-AAL*01_S2543 | IGKJ3-I3 | HQHNRYPHT | 9 | 4.4% | 289 glycan hole |
| RM19B | week 22 | rh1987 | Lambda | IGHV1-AAU*01_S5608 | IGHD6-11*01 | IGHJ5-2*01 | ARIPWRSAIAPGMRGNSFDV | 20 | 4.4% | IGLV2-ABJ*01_S3398 | IGLJ1-S1 | SSYAGSNSYI | 10 | 4.5% | base |
| RM19B1 | week 22 | rh1987 | Lambda | IGHV1-AAU*01_S5608 | IGHD6-11*01 | IGHJ5-2*01 | ARIMWRSAIAPGMRGNSMDV | 20 | 6.7% | IGLV2-ABJ*01_S3398 | IGLJ1-S1 | SSYGGSNTYI | 10 | 6.0% | base |
| RM19C | week 22 | rh1987 | Lambda | IGHV4-AGU*01_S6650 | IGHD2-8*01 | IGHJ5-2*01 | ARYCRGSICYGGGSLDV | 17 | 7.9% | IGLV2-ABE*01 | IGLJ3-S3 | SSYAGSNTLV | 10 | 1.8% | base |
| RM19C2 | week 25 | rh1987 | Lambda | IGHV4-AGU*01_S6650 | IGHD2-8*01 | IGHJ5-2*01 | ARYCSGSLCYGGGSLDV | 17 | 9.8% | IGLV2-ABE*01 | IGLJ3-S3 | SSYAGSNTLV | 10 | 4.3% | base |
| RM19C3 | week 25 | rh1987 | Lambda | IGHV4-AGU*01_S6650 | IGHD2-8*01 | IGHJ5-2*01 | ARYCSGALCYGGGSLDV | 17 | 7.3% | IGLV2-ABE*01 | IGLJ3-S3 | SSYAGSNTLL | 10 | 1.8% | base |
| RM19C4 | week 25 | rh1987 | Lambda | IGHV4-AGU*01_S6650 | IGHD2-8*01 | IGHJ5-2*01 | ARYCRGSICYGGGSLDV | 17 | 10.2% | IGLV2-ABE*01 | IGLJ3-S3 | SSYAGNNILF | 10 | 2.7% | base* |
| RM19E | week 22 | rh1987 | Lambda | LJI.Rh_IGHV3.122_S8388 | IGHD2-35*01 | LJI.Rh_IGHJ5.4 | VRVVVSATRGDHFDV | 15 | 6.3% | IGLV8-ABK*01 | IGLJ6-S6 | LIYMGSGISV | 10 | 6.2% | base |
| RM19F | week 22 | rh1987 | Lambda | LJI.Rh_IGHV4.79.a | IGHD2-35*01 | IGHJ4*01 | ARFLVVSAIKAATVPLEY | 18 | 3.9% | IGLV2-ABE*01 | IGLJ3-S3 | SSYAGSNTYVL | 11 | 0.9% | base |
| RM19F1 | week 22 | rh1987 | Lambda | LJI.Rh_IGHV4.79.a | IGHD2-35*01 | IGHJ4*01 | ARFLVVSAIKAATVPLEY | 18 | 5.1% | IGLV2-ABE*01 | IGLJ3-S3 | SSYAGSNTYVL | 11 | 0.9% | n.d. |
| RM19G | week 22 | rh1987 | Lambda | IGHV1-AAU*02_S1575 | IGHD6-6*01 | IGHJ4*01 | ARDRLAAAGTEHFDY | 15 | 4.2% | LJI.Rh_IGLV2.33.a | IGLJ3*01 | CSYLSGSTWV | 10 | 3.0% | base |
| RM19K | week 25 | rh1987 | Lambda | IGHV1-AAU*01_S5608 | IGHD3-26*01 | IGHJ5-2*01 | ARGLDYYGARQLGNSLHV | 18 | 6.1% | LJI.Rh_IGLV2.29 | IGLJ3*01 | YSYAGSNTWV | 10 | 3.0% | base |
| RM19L | week 25 | rh1987 | Lambda | LJI.Rh_IGHV4.67.a | IGHD2-35*01 | IGHJ3*01 | ARAPSYGGTSWAFSEGFDF | 19 | 6.2% | IGLV11-AAY*01 | IGLJ2-S2 | QMYGSSGAL | 9 | 3.8% | base |
| RM19M | week 25 | rh1987 | Lambda | IGHV4-AGU*01 | IGHD3-9*01 | LJI.Rh_IGHJ5.4 | ARDGYENDYGFYHPVRHRFDV | 21 | 8.4% | LJI.Rh_IGLV2.112 | IGLJ3*01 | FSYTTNKTWV | 10 | 10.6% | base |
| RM19O | week 25 | rh1987 | Lambda | IGHV4-AGR*01 | IGHD3-26*01 | IGHJ4*01 | AAGGGYYGNHFDF | 13 | 5.9% | IGLV2-ABE*01_S2946 | IGLJ1*01 | SSYAGSSTHLF | 11 | 3.1% | base |
| RM20E | week 53 | rh2011 | Kappa | IGHV5-ABI*01_S2502 | IGHD1-7*01 | IGHJ5-1*01 | VMWVYILTTGNIWVDV | 16 | 6.4% | LJI.Rh_IGKV2.71 | IGKJ2-I2 | GQITDFPYS | 9 | 3.6% | N611/FP |
| RM20E1 | week 53 | rh2011 | Kappa | IGHV5-ABI*01_S2502 | IGHD1-7*01 | IGHJ5-1*01 | VMWVYILTTGNIWVDV | 16 | 5.4% | LJI.Rh_IGKV2.71 | IGKJ2-I2 | GQITDFPYS | 9 | 3.6% | N611/FP |
| RM20E2 | week 53 | rh2011 | Kappa | IGHV5-ABI*01_S2502 | IGHD1-7*01 | IGHJ5-1*01 | VMWVYILTTGNIWVDV | 16 | 5.1% | LJI.Rh_IGKV2.71 | IGKJ2-I2 | GQITDFPYS | 9 | 4.5% | N611/FP* |
| RM20E3 | week 53 | rh2011 | Kappa | IGHV5-ABI*01_S2502 | IGHD1-7*01 | IGHJ5-1*01 | VMWVYILTTGNIWVDV | 16 | 5.4% | LJI.Rh_IGKV2.71 | IGKJ2-I2 | GQGTDFPYS | 9 | 3.0% | N611/FP* |
| RM20F | week 53 | rh2011 | Kappa | IGHV3-AFY*10 | IGHD3-14*01 | IGHJ2-I1 | ARGGKPIYYSGGYPSWYFDL | 20 | 7.3% | IGKV6-ABK*01 | IGKJ3-I3 | QQTNSFPCT | 9 | 5.3% | gp120/gp41 interface |
| RM20H | week 53 | rh2011 | Kappa | LJI.Rh_IGHV3.76.a_S4190 | IGHD2-13*01 | IGHJ5-1*01 | AKDWVGDYYTGTYPGFWFDV | 20 | 6.0% | IGKV3-ADW*01 | IGKJ4-I4 | QQNSNWPLT | 9 | 4.0% | gp120/gp41 interface |
| RM20I | week 53 | rh2011 | Kappa | IGHV4-ACQ*01_S4946 | IGHD3-3*01 | IGHJ4*01 | ARRVSRLDFGGAFDC | 15 | 6.2% | LJI.Rh_IGKV2.71.a | IGKJ4-I4 | GQGTHWPLT | 9 | 3.9% | n.d. |
| RM20J | week 53 | rh2011 | Kappa | IGHV4-AGU*01_S6650 | IGHD1-1*01 | IGHJ4*01 | ARWSTADFDY | 10 | 8.2% | IGKV1-AAL*01_S2543 | IGKJ3-I3 | QQHNNYPLT | 9 | 5.1% | 289 glycan hole |
| RM20A | week 22 | rh2011 | Lambda | IGHV3-AFE*01_S3736 | IGHD6-29*01 | IGHJ4*01 | AKGGMSSAWQSSKYYFDF | 18 | 5.3% | IGLV2-ABE*01_S2946 | IGLJ1-S1 | SSYAGSKTFYI | 11 | 4.2% | base* |
| RM20A1 | week 22 | rh2011 | Lambda | IGHV3-AFE*01_S3736 | IGHD6-29*01 | IGHJ4*01 | ARGGMSSAWQSSKYYFDQ | 18 | 8.5% | IGLV2-ABE*01_S2946 | IGLJ1-S1 | SSYAGRNTFYV | 11 | 6.4% | base* |
| RM20A2 | week 25 | rh2011 | Lambda | IGHV3-AFE*01_S3736 | IGHD6-29*01 | IGHJ4*01 | ARGGMSSAWQSSKYYFDQ | 18 | 9.1% | IGLV2-ABE*01_S2946 | IGLJ1-S1 | SSYAGRNTFYV | 11 | 4.6% | base |
| RM20A3 | week 53 | rh2011 | Lambda | IGHV3-AFE*01_S3736 | IGHD6-29*01 | IGHJ4*01 | ATGGMSSALQSSKYYFDF | 18 | 9.1% | IGLV2-ABE*01_S2946 | IGLJ1-S1 | SAYAGRQTFYI | 11 | 4.8% | base |
| RM20B | week 25 | rh2011 | Lambda | IGHV4-S11 | IGHD2-35*01 | IGHJ5-1*01 | ARLLVSAIRWEDRFDV | 16 | 3.0% | LJI.Rh_IGLV2.18 | IGLJ1-S1 | SSLGGSGTYI | 10 | 5.1% | base |
| RM20B1 | week 25 | rh2011 | Lambda | IGHV4-S11 | IGHD2-35*01 | IGHJ5-1*01 | ARLLVSAIRWEDRFDV | 16 | 3.8% | LJI.Rh_IGLV2.18 | IGLJ1-S1 | SSFAGGGTYI | 10 | 4.7% | base |
| RM20C | week 25 | rh2011 | Lambda | IGHV3-AEW*01 | IGHD3-9*01 | IGHJ4*01 | TRVAYEHDYGYYYKGYYFDF | 20 | 4.4% | IGLV1-ACV*01 | IGLJ6-S6 | QSYDSSLSAHV | 11 | 4.5% | base |
| RM20D | week 25 | rh2011 | Lambda | IGHV3-AFY*05 | IGHD4-15*01 | IGHJ4*01 | ARDPSRYGNYPDN | 13 | 3.6% | IGLV4-ACF*02 | IGLJ3-S3 | QTWTAGIVS | 9 | 2.1% | n.d. |
| RM20G | week 53 | rh2011 | Lambda | IGHV3-AEW*01 | LJI.Rh_IGHD5.29 | IGHJ6*01 | TRMRFASAQEPVYYGMD | 17 | 6.7% | IGLV1-S16_S2133 | IGLJ1-S1 | AAWDDSLGGYI | 11 | 3.3% | base |

***Inferred based on clonal relationship to mAbs with epitopes mapped by ns-EM.**
**Clonal families are highlighted with individual colors.**

**Supplementary Table 3. BLI Binding Kinetics**

| mAb | KD (M) | kon(1/Ms) | kdis(1/s) |
|---|---|---|---|
| RM19A | n.d. | n.d. | n.d. |
| RM19A1 | 3.94E-10 | 1.41E+04 | 5.57E-06 |
| RM19A2 | n.d. | n.d. | n.d. |
| RM19A3 | n.d. | n.d. | n.d. |
| RM19B | 9.87E-10 | 3.12E+05 | 3.08E-04 |
| RM19B1 | 7.00E-10 | 4.01E+05 | 2.80E-04 |
| RM19C | 1.50E-09 | 2.66E+05 | 4.00E-04 |
| RM19C2 | <1.0E-12 | 1.12E+05 | <1.0E-07 |
| RM19C3 | 6.80E-11 | 9.95E+04 | 6.76E-06 |
| RM19C4 | n.d. | n.d. | n.d. |
| RM19D | 1.57E-07 | 1.68E+03 | 2.63E-04 |
| RM19E | 1.28E-10 | 2.80E+05 | 3.60E-05 |
| RM19F | 4.98E-09 | 6.30E+05 | 3.14E-03 |
| RM19F1 | 2.88E-08 | 7.98E+04 | 2.30E-03 |
| RM19G | 3.95E-07 | 2.67E+03 | 1.05E-03 |
| RM19J | 1.95E-08 | 3.98E+03 | 7.77E-05 |
| RM19K | 8.55E-08 | 3.72E+03 | 3.18E-04 |
| RM19L | 6.03E-10 | 1.02E+05 | 6.15E-05 |
| RM19M | 1.03E-08 | 2.27E+04 | 2.34E-04 |
| RM19N | 1.46E-08 | 3.66E+05 | 5.33E-03 |
| RM19O | 1.87E-08 | 3.82E+04 | 7.14E-04 |
| RM19P | 2.09E-09 | 8.26E+03 | 1.73E-05 |
| RM19R | 5.52E-10 | 1.12E+05 | 6.18E-05 |
| RM19S | 1.47E-07 | 7.16E+03 | 1.05E-03 |
| RM19T | 8.45E-10 | 2.28E+04 | 1.93E-05 |
| RM20A | n.d. | n.d. | n.d. |
| RM20A1 | n.d. | n.d. | n.d. |
| RM20A2 | <1.0E-12 | 8.95E+05 | <1.0E-07 |
| RM20A3 | <1.0E-12 | 7.82E+05 | <1.0E-07 |
| RM20B | 3.92E-08 | 1.67E+04 | 6.56E-04 |
| RM20B1 | 1.85E-09 | 3.48E+05 | 6.44E-04 |
| RM20C | 2.55E-09 | 2.67E+04 | 6.79E-05 |
| RM20D | n.d. | n.d. | n.d. |
| RM20E | n.d. | n.d. | n.d. |
| RM20E1 | 5.85E-10 | 5.75E+04 | 3.36E-05 |
| RM20E2 | n.d. | n.d. | n.d. |
| RM20E3 | n.d. | n.d. | n.d. |
| RM20F | 1.60E-08 | 8.11E+03 | 1.30E-04 |
| RM20G | 8.01E-11 | 5.71E+05 | 4.58E-05 |
| RM20H | 1.61E-08 | 1.37E+04 | 2.21E-04 |
| RM20I | 1.27E-07 | 2.53E+03 | 3.21E-04 |
| RM20J | <1.0E-12 | 4.47E+04 | <1.0E-07 |

**Supplementary Table 4. EM Data Collection and Map/Model Refinement Parameters**

| Complex | BG505.v4.1 + RM20F | BG505.v5.2 + RM20J | BG505.v5.2 + RM20E1 + PGT122 |
|---|---|---|---|
| Microscope | Titan Krios | Talos Arctica | Talos Arctica |
| Voltage, kV | 300 | 200 | 200 |
| Detector | Gatan K2 Sumit | Gatan K2 Sumit | Gatan K2 Sumit |
| Recording Mode | Counting | Counting | Counting |
| Magnification | 29,000 | 36,000 | 36,000 |
| Moive micrograph pixel size, Å | 1.03 | 1.15 | 1.15 |
| Dose rate, $e^-$/[(camera pixel)*s] | 8.833 | 6.004 | 5.656 |
| No. of frames per moive micrograph | 24 | 44 | 48 |
| Frame exposure time, ms | 250 | 250 | 250 |
| Movie micrograph exposure time, s | 6 | 11 | 12 |
| Total dose, $e^-/Å^2$ | 50.0 | 49.9 | 51.3 |
| Defocus range, µm | 1.3 to 2.8 | 1.0 to 3.5 | 1.0 to 2.5 |
| No. of movie micrographs | 1055 | 553 | 579 |
| No. of molecular projection images in map | 91212 | 26327 | 17010 |
| Symmetry | C3 | C3 | C1 |
| Map resolution (FSC 0.143) | 4.25 | 3.88 | 4.42 |
| Map sharpening B-factor, $Å^2$ | -174.7 | -112.6 | -81.2 |
| No. of atoms in deposited model | 19962 | 19785 | 21768 |
| MolProbity score | 1.16 | 1.38 | 0.97 |
| $C\beta$ Outliers (%) | 0.00 | 0.00 | 0.00 |
| Rotamer Outliers (%) | 0.15 | 0.72 | 0.13 |
| Rama Outliers (%) | 0.65 | 1.29 | 0.20 |
| Clashscore | 1.75 | 2.12 | 0.97 |
| EMRinger score | 2.02 | 2.71 | 1.82 |
| Privateer | pass | pass | pass |
| EMDB | EMD-21246 | EMD-21257 | EMD-21232 |
| PDB ID | 6VN0 | 6VO1 | 6VLR |

**Supplementary Table 5.** X-ray data collection and refinement statistics

| **Data collection** | **RM20F (PDB:6VSR)** | **RM20J (PDB:6VOS)** | **RM20E1 (PDB:6VOR)** |
|---|---|---|---|
| Beamline | SSRL 12-2 | APS 23ID-B | SSRL 12-2 |
| Wavelength (Å) | 0.97946 | 1.03322 | 0.97946 |
| Space group | $P\ 2_1\ 2_1\ 2_1$ | $P\ 4_3\ 2_1\ 2$ | P 1 |
| Unit cell parameters | a=61.5, b=75.6, c=115.1, α= β=γ=90 | a=b=112.3, c=141.0, α=β=γ=90 | a=57.4, b=57.7, c=92.7, α=98.7, β=94.1, γ=97.8 |
| Resolution (Å) | 36.7-2.20 (2.24-2.20)[a] | 50.0-2.30 (2.34-2.30)[a] | 50.0-1.85 (1.90-1.85)[a] |
| Unique Reflections | 28,053 (1,364)[a] | 40,057 (1,946)[a] | 142,088 (2,413)[a] |
| Redundancy | 4.0 (4.0)[a] | 17.5 (11.2)[a] | 2.7 (1.3)[a] |
| Completeness (%) | 98.6 (99.4)[a] | 100 (99.6)[a] | 85.4 (32.5)[a] |
| $\langle I/\sigma_I \rangle$ | 7.7 (2.0)[a] | 28.0 (1.0)[a] | 18.8 (1.2)[a] |
| $R_{sym}$[b] (%) | 19.7 (83.0)[a] | 12.9 (>100)[a] | 10.9 (69.1)[a] |
| $R_{pim}$[b] (%) | 11.0 (46.4)[a] | 3.0 (50.3)[a] | 4.9 (51.7)[a] |
| $CC_{1/2}$[c] (%) | 83.9 (54.0)[a] | 89.7 (44.1)[a] | 91.6 (56.0)[a] |
| **Refinement statistics** | | | |
| Reflections (work) | 26,622 | 39,953 | 75,965 |
| Reflections (test) | 1,374 | 1,999 | 3,285 |
| $R_{cryst}$[d] / $R_{free}$[e] (%) | 19.7/24.1 | 18.1/21.1 | 22.4/25.2 |
| No. of atoms | | | |
| Protein | 3,328 | 3,604 | 6,731 |
| Water | 288 | 224 | 320 |
| Average *B*-value (Å$^2$) | | | |
| Protein | 25 | 62 | 39 |
| Water | 34 | 68 | 40 |
| Wilson *B*-value (Å$^2$) | 21 | 57 | 30 |
| **RMSD from ideal geometry** | | | |
| Bond length (Å) | 0.002 | 0.007 | 0.008 |
| Bond angle (°) | 0.55 | 0.95 | 0.99 |
| **Ramachandran statistics (%)** | | | |
| Favored | 97.5 | 96.1 | 96.3 |
| Outliers | 0.0 | 0.2 | 0.1 |

[a] Numbers in parentheses refer to the highest resolution shell.

[b] $R_{sym} = \Sigma_{hkl} \Sigma_i | I_{hkl,i} - \langle I_{hkl} \rangle | / \Sigma_{hkl} \Sigma_i I_{hkl,i}$ and $R_{pim} = \Sigma_{hkl} (1/(n-1))^{1/2} \Sigma_i | I_{hkl,i} - \langle I_{hkl} \rangle | / \Sigma_{hkl} \Sigma_i I_{hkl,i}$, where $I_{hkl,i}$ is the scaled intensity of the i$^{th}$ measurement of reflection h, k, l, $\langle I_{hkl} \rangle$ is the average intensity for that reflection, and *n* is the redundancy.

[c] $CC_{1/2}$ = Pearson correlation coefficient between two random half datasets.

[d] $R_{cryst} = \Sigma_{hkl} | F_o - F_c | / \Sigma_{hkl} | F_o |$ x 100, where $F_o$ and $F_c$ are the observed and calculated structure factors, respectively.

[e] $R_{free}$ was calculated as for $R_{cryst}$, but on a test set comprising 5% of the data excluded from refinement.
